# Perineuronal Net and Inhibitory Synapse Remodeling on Striatal Fast-spiking Interneurons by Chronic Alcohol Exposure

**DOI:** 10.1101/2025.07.08.663744

**Authors:** Michael S. Patton, Samuel H. Sheats, Andreas B. Wulff, Paige N. McKeon, Jonathan W. VanRyzin, Mary H. Patton, Morgan Heckman, Allison N. Siclair, Phillip H. Iffland, Brian N. Mathur

## Abstract

Alcohol use disorder is characterized by persistent drinking in the face of negative consequences. Such inflexible drinking requires dorsolateral striatum fast-spiking interneurons, which comprise roughly 1% of all striatal neurons. How chronic ethanol exposure affects fast-spiking interneuron physiology is poorly understood. We discover in mice that chronic ethanol exposure induced a dramatic loss of GABAergic, but not glutamatergic, synapses onto dorsolateral striatum fast-spiking interneuron somata and proximal dendrites where perineuronal nets, a subdivision of the extracellular matrix, are enriched. We found that chronic ethanol exposure degraded these perineuronal nets and that enzymatically degrading perineuronal nets similarly reduced GABAergic transmission onto dorsolateral striatum fast-spiking interneurons. Modeling the effect of alcohol, we find that silencing extrinsic GABAergic projections to the dorsolateral striatum increased voluntary ethanol consumption. Taken together, these data suggest chronic alcohol exposure remodels perineuronal nets and inhibitory synapses on fast-spiking interneurons to facilitate alcohol drinking.

## Introduction

Inflexible drinking is a prominent feature of alcohol use disorder that is not adequately addressed by current therapies^1^. The dorsolateral striatum encodes inflexible actions that are thought to underlie inflexible drinking in alcohol use disorder^2–6^. Indeed, ethanol exposure promotes activity in the dorsolateral striatum and biases behavior from goal-directed to inflexible, stimulus-response-based strategies^7–12^. This includes inflexible consumption of both natural rewards^11^ and alcohol^12, 13^.

Parvalbumin-expressing fast-spiking interneurons (FSIs) are enriched in the dorsolateral striatum and provide feedforward inhibition onto striatal projection neurons^14–17^. Habitual responding for natural rewards requires FSIs^18^ and FSIs are necessary for the expression of organized, aversion resistant alcohol drinking behavior^13^. Thus, FSIs are positioned to shape dorsolateral striatal medium spiny neuron encoding of stereotyped behaviors ^19–22^. Exploring how alcohol may be facilitating this FSI-mediated process, several studies identified that acute alcohol targets FSIs ^23–26^. What is not known, however, is how chronic alcohol exposure, modeling the human experience in alcohol use disorder, affects FSI synaptic physiology and related behavior.

Dorsolateral striatum FSIs receive excitatory innervation primarily from the somatomotor cortices and receive inhibitory innervation primarily from the reticular thalamic nucleus (RTN) and the globus pallidus (GP) ^27–31^. Projections from the GP to the dorsolateral striatum are implicated in the transition from goal-directed to habitual behavior^32^ and acute alcohol potentiates GABAergic synaptic transmission at GP and RTN synapses onto FSIs^23^. This suggests that chronic alcohol exposure may result in lasting changes at these synapses to promote drinking. In the following work, we demonstrate that chronic ethanol vapor exposure reduced GABAergic, but not glutamatergic, synaptic transmission onto dorsolateral striatum FSIs from both the GP and RTN. This reduction in GABAergic transmission was mediated by a decrease in the number of GABAergic synapses, primarily on somata and proximal dendrites of FSIs where perineuronal nets (PNNs) uniquely surround FSIs. Following this observation, we discovered that chronic ethanol exposure reduced the number of PNN-positive FSIs as well as striatal PNN protein expression, while enzymatic digestion of PNNs reduced the number of GABAergic synapses on FSIs. Silencing inhibitory synaptic transmission to the dorsolateral striatum, mirroring what is observed following chronic ethanol exposure, increased voluntary ethanol consumption. These data suggest a dramatic remodeling of inhibitory control of FSIs in the dorsolateral striatum following chronic alcohol exposure contributes to drinking phenotypes.

## Methods

### Animals

All experiments were performed in accordance with NIH guidelines and were approved by the Institutional Animal Care and Use Committee of the University of Maryland Baltimore and the National Institute on Alcohol Abuse and Alcoholism. Randomly sampled 2-4 month old male and female wildtype C57BL/6J mice or mice expressing cre recombinase under the parvalbumin promoter^33^ crossed with a tdTomato reporter mouse line (PV-tdT) were housed with littermates (2-5 per cage) under a normal 12-hour light/dark cycle (lights on at 0900 hours, off at 2100 hours) with *ad libitum* access to food and water.

### Surgical procedures

All stereotaxic injections were performed under isoflurane anesthesia (5% induction; 2%–3% maintenance) and viruses were injected at a rate of 30nl/min with a 25G syringe (Hamilton Company). To isolate GABAergic projections from the globus pallidus (GP) or the reticular thalamic nucleus (RTN) to the dorsolateral striatum, PV-tdT mice on a C57BL/6J background^33, 34^ were stereotaxically injected with a virus expressing channelrhodopsin2 (ChR2; AAV5-hSyn-ChR2-eYFP) in the GP (AP −0.4 mm, ML ± 2.0 mm, DV −3.7 mm; 200 nl/side) or the RTN (AP −0.58 mm, ML ± 1.25 mm, DV −3.5 mm; 200 nl/side). To degrade PNNs, PV-tdT mice were stereotaxically injected with chondroitinase ABC (ChABC: 15 units/mL in PBS with 1% bovine serum albumin (BSA); Sigma) or vehicle unilaterally into the dorsolateral striatum (AP +0.60 mm, ML ± 2.25 mm, DV −3.0 mm; 500 nl/side). To label postsynaptic GABAergic density within PV cells, PV-tdT mice were stereotaxically injected with a fibronectin intrabody against gephyrin (AAV5-EF1A-DIO-Gephyrin.FingR-eGFP^35^) in the dorsolateral striatum (AP +0.60 mm, ML ± 2.25 mm, DV −3.0 mm; 500 nl/side). To silence inhibitory transmission from the GP and RTN to the dorsolateral striatum, C57BL/6J mice were stereotaxically injected with 250nl rgAAV-Cre-mCherry (Addgene # 55632) and 200 nl AAV5-FLEX-TeLC-GFP (UMB Vector Core, Addgene Plasmid # 135391) or AAV5-DIO-GFP (Addgene # 27056) in the GP and RTN.

### Chronic Intermittent Ethanol Exposure (CIE)

Randomly sampled male and female mice were placed in plexiglass inhalation chambers (60×36×60 cm)^36^ and exposed to ethanol vapor or air 16 hours/day, four days a week for a maximum of five weeks. After four days in the inhalation chambers, mice underwent a 72-hour forced abstinence period from ethanol. Vapor chamber ethanol concentrations were monitored daily, and air flow was adjusted to produce ethanol concentrations within 1.8-2.0 % ethanol content measured by a digital alcohol breath tester (FFtopu). These conditions produce stable blood ethanol concentrations in C57BL/6J mice ranging from 150-200 mg/dl^37^. The alcohol dehydrogenase inhibitor pyrazole (1 mmol/kg, i.p.) was injected into both control and ethanol vapor chamber mice during weeks 4-5 to stabilize blood ethanol concentrations.

### Immunohistochemistry

Mice were transcardially perfused with room-temperature 0.1 M Phosphate Buffered Saline (PBS), pH 7.3, followed by room-temperature 4% (w/v) paraformaldehyde in PBS. Brains were extracted and post-fixed with 4% paraformaldehyde in PBS at 4°C for 24-48h. Coronal sections (50 μm) were cut using a Leica VT1000S vibrating microtome. For PNN analysis, serial sections were stained with Wisteria floribunda agglutinin (WFA) conjugated to fluorescein (1:1,000; ThermoFisher # L32481). Fast-spiking interneurons were fluorescently labeled with an anti-PV antibody (mouse; 1:1,1000; Sigma # P3088). Presynaptic GABAergic terminals were fluorescently labeled with an anti-vesicular GABA transporter (VGAT) antibody (rabbit; 1:1,000; Synaptic Systems # 131002). Slices were incubated in secondary antibodies conjugated to Alexa 594 (Donkey; 1:1000) at room temp for 4 h.

### 3D neuronal reconstruction

Striatal fast-spiking interneurons were filled with biocytin (5%; Tocris # 3349) with a borosilicate glass pipette (3-5 MΩ resistance) under 40x immersion objective. Slices (250 µm) were fixed in 4% paraformaldehyde overnight at 4°C. The next day, slices were washed in 0.1 M PBS (3 x 20 min) and blocked with 1% BSA in PBS + 0.3% Triton X-100 (PBS-T) for 2 h at room temperature. Slices were incubated with Alexa Fluor 594-streptavidin (Invitrogen, # S32356) (1:1000, 1% BSA in 0.3% PBS-T) overnight at 4°C. The next day, slices were washed in PBS (3 x 20 min), mounted on slides, and cover-slipped using ProLong Diamond Antifade (Invitrogen # P36965). Neurons were reconstructed in Neurolucida 360 (version 2020.1.1) using semi-automated reconstruction with a directional kernels algorithm.

### Microscopy

For 3D neuronal reconstruction, confocal images were acquired with a Nikon W1 microscope equipped with a 561 nm laser. Neurons were imaged using a 40x oil (1.30 NA) objective with a lateral resolution of 0.160 µm per pixel and a 0.200 µm z-step. For PNN analysis and gephyrin-VGAT colocalization, confocal images were acquired with a Nikon A1 microscope equipped with a 488 and 561 nm lasers. Neurons were imaged using a 40x oil (1.40 NA) objective with a lateral resolution of 0.2 µm per pixel and a 0.20 µm z-step. Images were denoised with Nikon NIS-ElementsDenoise.ai software.

Surface models and filament models were generated using Imaris Cell Imaging Software (Oxford Instruments). Colocalized gephyrin-vGAT puncta were quantified using the dot-model function with masking application in Imaris. Fluorescent images for population PNN analysis were visualized on a Nikon Eclipse Ti2 fluorescent microscope and captured with a Nikon DS-Qi2 camera. PNNs surrounding FSIs were analyzed by a fluorescent intensity profile threshold, where enriched FSIs contained a signal of ≥ 1000 intensity units from background and undetected PNNs were classified by a signal intensity < 500 intensity units from background.

### Whole-cell patch-clamp electrophysiology

Mice were deeply anesthetized with isoflurane before rapid decapitation and brain removal. 250 µm coronal sections were collected in ice cold modified artificial cerebral spinal fluid (aCSF: 194 mM sucrose, 30 mM NaCl, 4.5 mM KCl, 1 mM MgCl_2_, 26 mM NaHCO_3_, 1.2 mM NaH_2_PO_4_, and 10 mM D-glucose bubbled with 95% oxygen, 5% carbon dioxide) using a Leica VT1200 vibrating microtome. Brain sections were then transferred to regular aCSF (124 mM NaCl, 4.5 mM KCl, 2 mM CaCl_2_, 1 mM MgCl_2_, 26 mM NaHCO_3_, 1.2 mM NaH_2_PO_4_, and 10 mM D-glucose, bubbled with 95% oxygen, 5% carbon dioxide) and incubated at 32°C for 30 min for recovery, followed by room temperature until recording. Slices were hemisected, placed into a recording chamber, and perfused with temperature-controlled aCSF (29-31°C) during the recording procedure (aCSF: 124 mM NaCL, 4.5 mM KCl, 1 mM MgCl_2_, 26 mM NaHCO_3_, 1.2 mM NaH_2_PO_4_, 10 mM D-glucose, 2 mM CaCl_2_, bubbled with 95% oxygen, 5% carbon dioxide). Dorsolateral striatum FSIs were fluorescently visualized using Eclipse FN1 Nikon microscope and Intenslight C-HGFIE (Nikon). Images were digitally rendered using a Hamamatsu ORCA-ER digital camera and associated HC image live 4.8.0 software. Striatal FSIs were voltage-clamped at −60 mV using a MultiClamp 700B Amplifier (Molecular Devices). Electrically-evoked postsynaptic currents were generated using a constant current isolated stimulator (Digitimer Ltd model DS3) and a concentric bipolar stimulating electrode (World Precision Instruments) located approximately 150 µm from the recording electrode. Optogenetically-evoked currents were generated from an LED driver (Thorlabs, LEDD1B T-Cube) and a 470 nm LED delivering blue light (2 - 4ms pulse duration) through an optical fiber located approximately 150µm from the recording electrode. Excitatory postsynaptic currents (EPSCs) were pharmacologically isolated by adding picrotoxin (50 µM) to the recording aCSF. EPSCs were recorded using a borosilicate glass pipette (3-5 MΩ resistance) filled with cesium-based internal solution (120 mM CsMeSO_3_, 10 mM HEPES, 5 mM NaCl, 10 mM TEA-Cl, 1.1 mM EGTA, 0.3 mM Na-GTP, 5 mM QX-314, 4 mM Mg-ATP). Evoked inhibitory postsynaptic currents (IPSCs) and spontaneous IPSCs (sIPSCs) were pharmacologically isolated by adding DL-AP5 (50 µM) and CNQX (5 µM) to the recording aCSF. IPSCs were recording using a borosilicate glass pipette (3-5 MΩ resistance) filled with a high chloride-based internal solution (150 mM CsCl, 10 mM HEPES, 2 mM MgCl_2_, 0.3 mM Na-GTP, 5 mM QX-314, 3 mM Mg-ATP, and 0.2 mM BAPTA). For optically-evoked asynchronous currents, calcium was replaced with strontium (2 mM) in the recording aCSF. To photo-uncage GABA, a 2 ms pulse of blue light (470 nm) was delivered to an acute slice preparation bath perfused with 250 µM ruthenium-bipyridine-triphenylphosphine caged GABA^38^ (RuBi-GABA) (Tocris # 4709). Signals were filtered at 2k Hz, digitized at 10 kHz and acquired using Clampex 10.4.1.4 software (Molecular Devices). Data were analyzed offline using Clampfit software (Molecular Devices) and Mini Analysis Program (Synaptosoft).

### Western blot

Following CIE, mice were sacrificed by rapid decapitation and brains were immediately removed for tissue collection. The dorsal striatum (AP: +1.3 - 0.3 mm from bregma) was dissected from a 1 mm coronal section cut from a mouse brain mold. Left and right hemispheres were pooled and flash frozen in crushed dry ice. Tissue was stored at −80°C until further processing. Tissue lysates were prepared in RIPA lysis buffer (Milllipore # 20-188) containing protease (Promega G6521) and phosphatase inhibitors (Sigma # 4906845001). Protein content was normalized via BCA assay and 10µg of protein was loaded onto NuPAGE Novex Tris-Acetate mini protein gels (Novex # EA0375) and run with Tris-Acetate buffer (Novex # LA0041). Gels were electroblotted onto nitrocellulose membrane (0.45 µ pore size) using XCell II Blot Module (ThermoFisher). Membranes were blocked in Intercept blocking buffer (1:2 in TBS; Li-Cor) for 2h at room temperature before incubation overnight 4 °C in primary antibody: HAPLN1 (Goat; 1:1,000; R&D Systems # AF2608) or Aggrecan (Rabbit; 1:1,000; Millipore # AB1031). Membranes were washed three times for 10 min in TBS-T (0.1% Tween 20) before incubation in secondary antibodies: Donkey anti-rabbit (IRDye 926; 1:10,000; Li-Cor or Donkey anti-goat (IRDye 800; 1:10,000; Li-Cor). Images were captured using an Odyssey CLX (Li-Cor). Protein signal was normalized with a total protein stain (Revert total protein stain; Li-Cor) and data are represented as % control.

### Drinking in the Dark (DID)

The DID protocol was adapted from Rhodes et al.^39^ following our previously established methods^13^. Briefly, mice were randomly assigned to sucrose or ethanol drinking groups. Mice were given two hours of access to the drinking bottle in the first three days, and on day four, mice were given four hours of access. Following five days of drinking, mice underwent two days of forced abstinence, and this cycle repeated for two consecutive weeks. Bottle weights were recorded to determine total consumption and adjusted to body weight (g/kg).

### Statistical analysis

All statistical analyses were performed in GraphPad Prism 9.0. Data are represented as mean ± SEM. Comparisons between two groups were analyzed with a Student’s t-test while comparisons of three or more groups were analyzed with two-way ANOVA. Multiple comparisons were corrected with Sidak’s test or Dunnett’s test when comparing multiple time points to a single baseline. For the Sholl analysis, a mixed-effect model was performed^40^. All N are specifically defined in the legend of each figure. For data from CIE mice, N is defined as an individual animal. This N is derived from an average of 2-3 cell recordings per animal or an average fluorescent quantification of 5-10 cells per animal. For ChABC treatment, N is defined as an individual animal, with 2-3 cell recordings per animal. Both male and female mice were used in this study and data is pooled by sex.

## Results

We first performed whole-cell patch clamp recordings from dorsolateral striatum FSIs to determine if chronic intermittent ethanol (CIE) modulated synaptic strength onto FSIs (**Figure 1A,B**). Recording 4-7 days into abstinence, we found that CIE dramatically reduced electrically-evoked inhibitory transmission onto dorsolateral striatum FSIs (F_(1,10)_ = 19.59, p = 0.001; two-way repeated measures ANOVA; **Figure 1C**). However, we found no difference in the amplitude of electrically-evoked excitatory synaptic currents across increasing stimulus intensities (F_(1,10)_ = 0.058, p = 0.82; two-way repeated measures ANOVA; **Figure 1D**). Following these results, we recorded spontaneous inhibitory postsynaptic current (sIPSC) events onto dorsolateral striatum FSIs to determine if CIE-induced depression of these synapses was expressed pre- or post-synaptically. FSIs from CIE-exposed mice exhibited less frequent sIPSCs (increased inter-event interval, IEI) compared to FSIs from control mice (t_(18)_ = 3.12, p < 0.001; Student’s t-test; **Figure 1E,F**) but there was no change in the amplitude of sIPSCs (t_(18)_ = 1.47, p = 0.16; Student’s t-test; **Figure 1G).** In light of the significant change in inhibitory synaptic transmission, we assessed whether CIE impacted the cellular morphology of dorsolateral striatum FSIs. Sholl analysis revealed no statistically significant differences in dendritic complexity between control and CIE mice (F_(1,18)_ = 0.037, p = 0.85; Mixed-effect model^40^ **Figure 1H**).

**Figure 1.**
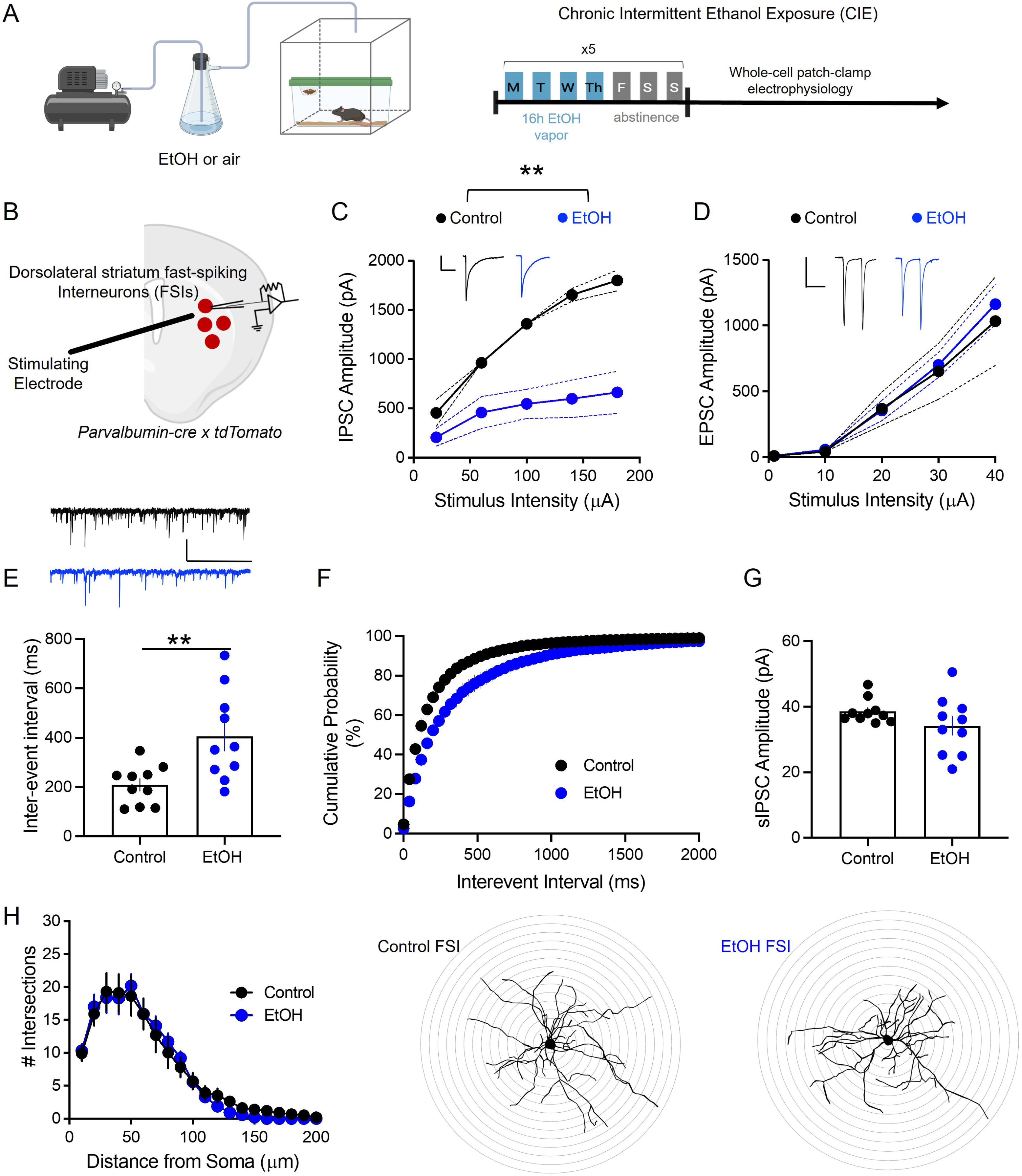
Chronic intermittent ethanol (CIE) vapor exposure specifically reduced inhibitory synaptic transmission onto dorsolateral striatum fast-spiking interneurons (FSIs). **A**) schematic and timeline of CIE vapor exposure paradigm. **B**) Schematic of whole-cell patch clamp recording from an FSI. **C**) CIE dramatically reduced inhibitory synaptic transmission onto FSIs (N=6 mice per group, 3M 3F). Inset: representative electrically evoked inhibitory postsynaptic currents (eIPSCs) from control (black) and CIE (blue) treated mice (scale bars 500pA, 100ms). **D**) However, we observed no change in electrically-evoked excitatory postsynaptic currents (eEPSCs) onto FSIs (N=6 mice per group, 3M 3F). **E, F**) Spontaneous inhibitory synaptic transmission onto dorsal striatum FSIs (N=10 mice per group, 5M 5F). Inset: representative traces from control (black) and CIE (blue) treated mice (scale bar 100pA, 2s). CIE reduced the frequency of spontaneous inhibitory events (**E, F**) but did not impact event amplitude (**G**). **H**) CIE did not impact FSI dendrite branch complexity (N=10 cells, 5 mice per group, 3M 2F). Representative dye-filled FSIs from control (left) and CIE (right) treated mice (Sholl ring radius = 10µm increments). All data represented as mean + SEM. * P<0.01. Portions of this figure were created with BioRender.com.

Considering that the data thus far suggest a suppression of presynaptic GABA release, we next examined the two major extrinsic sources of GABA to the dorsal striatum, the globus pallidus (GP) and the reticular thalamic nucleus (RTN), which synapse onto striatal FSIs^27, 29^. To investigate which distinct GABAergic projections were specifically impacted by CIE, we virally expressed channelrhodopsin (ChR2) into the GP (**Figure 2A**) or the RTN (**Figure 2F**) and recorded optically-evoked IPSCs onto dorsolateral striatum FSIs. GABAergic transmission from the GP was dramatically reduced by CIE (F_(1,70)_ = 42.45, p < 0.0001; two-way repeated measures ANOVA; **Figure 2B**), but there was no change in the paired-pulse ratio (PPR) from the GP (t_(14)_ = 1.07, p = 0.30, Student’s t-test; **Figure 2C**). Since we found a decrease in the frequency of sIPSCs (**Figure 1E, F**) but saw no difference in PPR (**Figure 2C**), we next conducted optically-evoked recordings where we replaced calcium with strontium in the external, artificial cerebrospinal fluid (aCSF) recording solution to elicit asynchronous synaptic release events from the GP. CIE reduced the frequency of asynchronous release events from the GP (t_(10)_ = 4.14, p < 0.005; Student’s t-test; **Figure 2D**) but had no impact on event amplitude (t_(10)_ = 0.09, p = 0.93; Student’s t-test; **Figure 2E**). Similar to GP inputs, CIE reduced the amplitude of optically-evoked IPSCs from the RTN onto dorsolateral striatum FSIs (F_(1,14)_ = 10.36, p < 0.01; two-way repeated measures ANOVA; **Figure 2G**). Additionally, CIE did not impact PPR from the RTN (t_(14)_ = 0.58, p = 0.57; Student’s t-test; **Figure 2H**). CIE reduced the frequency of optically-evoked strontium-enabled asynchronous inhibitory synaptic release events from the RTN (t_(10)_ = 6.20, p < 0.001; Student’s t-test; **Figure 2I**) but had no impact on event amplitude (t_(10)_ = 0.41, p = 0.69; Student’s t-test; **Figure 2J**).

**Figure 2.**
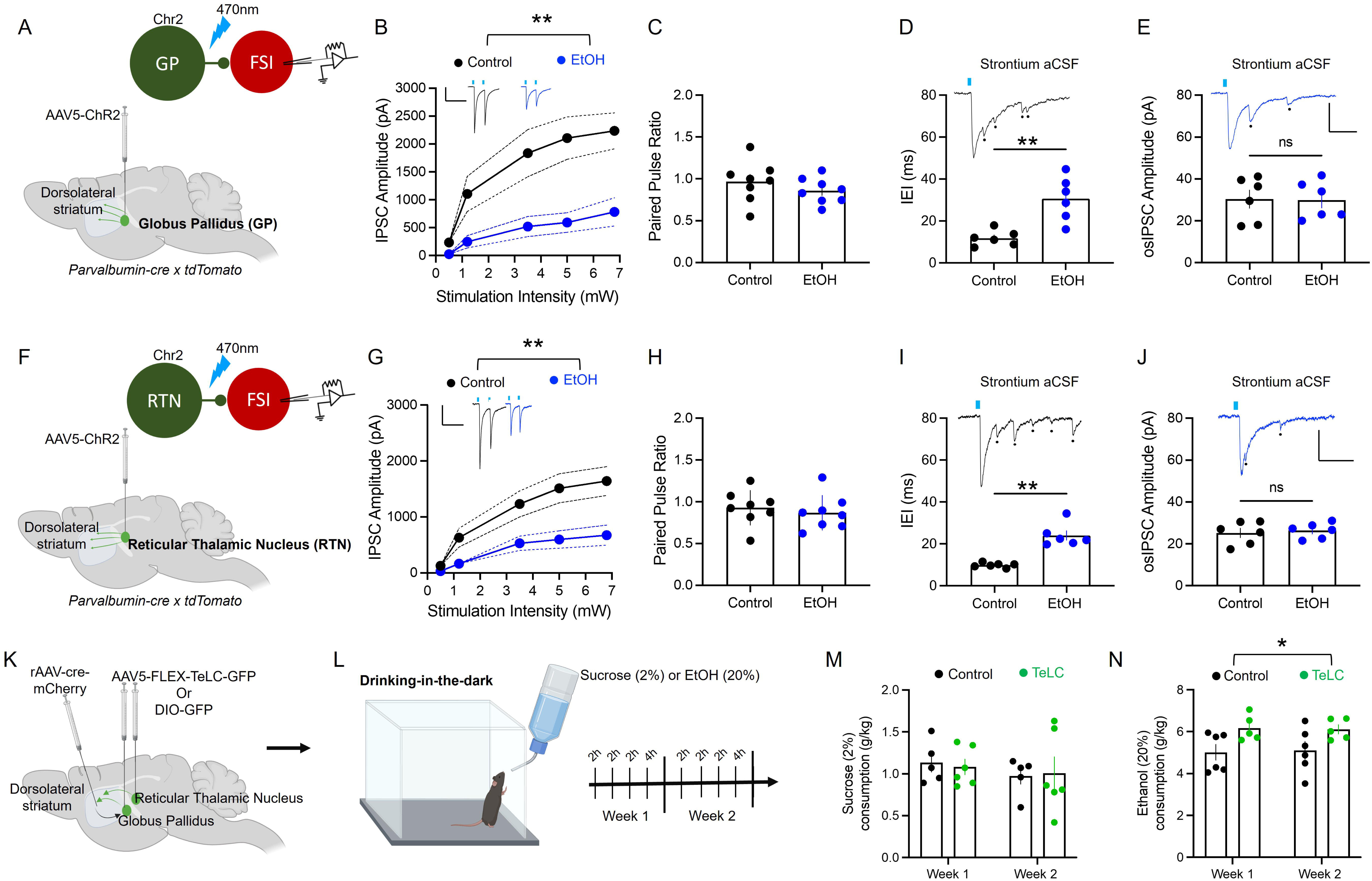
CIE reduced inhibitory synaptic transmission from the globus pallidus (GP) and the reticular thalamic nucleus (RTN) and silencing inhibitory transmission from these inputs increased ethanol consumption. **A**) Schematic of injection of a virus expressing channelrhodopsin (ChR2) into the globus pallidus. **B**) CIE reduced optically-evoked inhibitory currents from the GP onto FSIs but (**C**) had no impact on release probability (N=8 mice per group, 4M 4F) Inset: representative traces from control (black) and CIE (blue) treated mice (scale bar 500 pA, 100 ms). CIE reduced the (**D**) frequency but not the (**E**) amplitude of asynchronous inhibitory current events from the GP. Inset: representative traces from control (black) and CIE (blue) treated mice (scale bar 200 pA, 10 ms). **F**) Schematic of injection of virus expressing ChR2 into the RTN. **G**) CIE reduced optically-evoked inhibitory currents from the RTN but (**H**) had no impact on release probability (N=8 mice per group, 4M 4F; scale bar 500pa, 100ms). **I**) CIE reduced the frequency of asynchronous inhibitory current events but (**J**) did not impact asynchronous current event amplitude (N= 6 mice per group, 3M 3F; scale bar 200 pA, 20 ms). **K**) Schematic of injection of cre-dependent virus expressing tetanus light chain toxin (TeLC) into the GP and RTN and retrograde AAV expressing cre-recombinase in the dorsal striatum. **L**) Experimental timeline of Drinking in the Dark (DID) paradigm with schematic of mouse drinking ethanol (20%) or sucrose (2%). **M**) TelC silencing of inhibitory transmission from the GP and RTN did not impact sucrose consumption. **N**) However, TelC silencing of inhibitory transmission increased voluntary ethanol consumption. All data represented as mean + SEM. *P < 0.05, **P < .01. Portions of this figure were created with BioRender.com.

Next, we investigated whether virally silencing inhibitory projections to the dorsolateral striatum, mirroring what is observed following CIE, affects voluntary ethanol consumption in C57BL/6J mice. We microinjected a cre-dependent tetanus light chain (TeLC) into the GP and RTN and a retrograde AAV expressing cre-recombinase into the dorsolateral striatum (**Figure 2K**) to silence inhibitory synaptic transmission from the GP and RTN to the dorsolateral striatum synaptic transmission^41^. Following a 4-week recovery period, mice underwent a 2-week Drinking-in-the-dark (DID) paradigm (**Figure 2L)**. Silencing inhibitory projections from GP and RTN to the dorsolateral striatum did not affect sucrose drinking (**Figure 2M**) but increased voluntary ethanol consumption (F_(1,9)_=5.56, p<0.05; two-way repeated measures ANOVA; **Figure 2N**).

To examine the mechanism underlying the CIE-induced suppression of GABAergic transmission onto FSIs, we virally expressed a GFP intrabody against the postsynaptic GABAergic synapse marker, gephyrin, into dorsolateral striatum FSIs (**Figure 3A**) and immunostained for the presynaptic GABAergic marker vesicular GABA transporter (VGAT). We quantified colocalized VGAT- and gephyrin-positive puncta to identify putative GABAergic synapses (**Figure 3B**). CIE mice exhibited fewer GABAergic synapses onto dorsolateral striatum FSIs compared to control mice (**Figure 3B-E**). Notably, the loss of synapses was more pronounced on the somata (t_(14)_ = 4.491, p < 0.001; Student’s t-test; **Figure 3C**) and proximal branches of FSIs (t_(14)_ = 6.58, p < 0.0001; Student’s t-test; **Figure 3D**), but still present on distal processes (t_(14)_ = 2.16, p < 0.05; Student’s t-test; **Figure 3E**). To further investigate whether CIE resulted in a postsynaptic silencing of inhibitory synapses or merely a loss of presynaptic elements that leave postsynaptic sites intact (and unsilenced), we photo-uncaged RuBI-GABA in a slice preparation and recorded from FSIs using whole-cell patch-clamp electrophysiology in voltage clamp mode (**Figure 3F**). We found that uncaging IPSCs (uIPSCs) onto FSIs from control and CIE mice were not significantly different, suggesting that postsynaptic inhibitory synapses are not silenced (F_(1,14)_ = 0.19, p = 0.67; **Figure 3G,H**).

**Figure 3.**
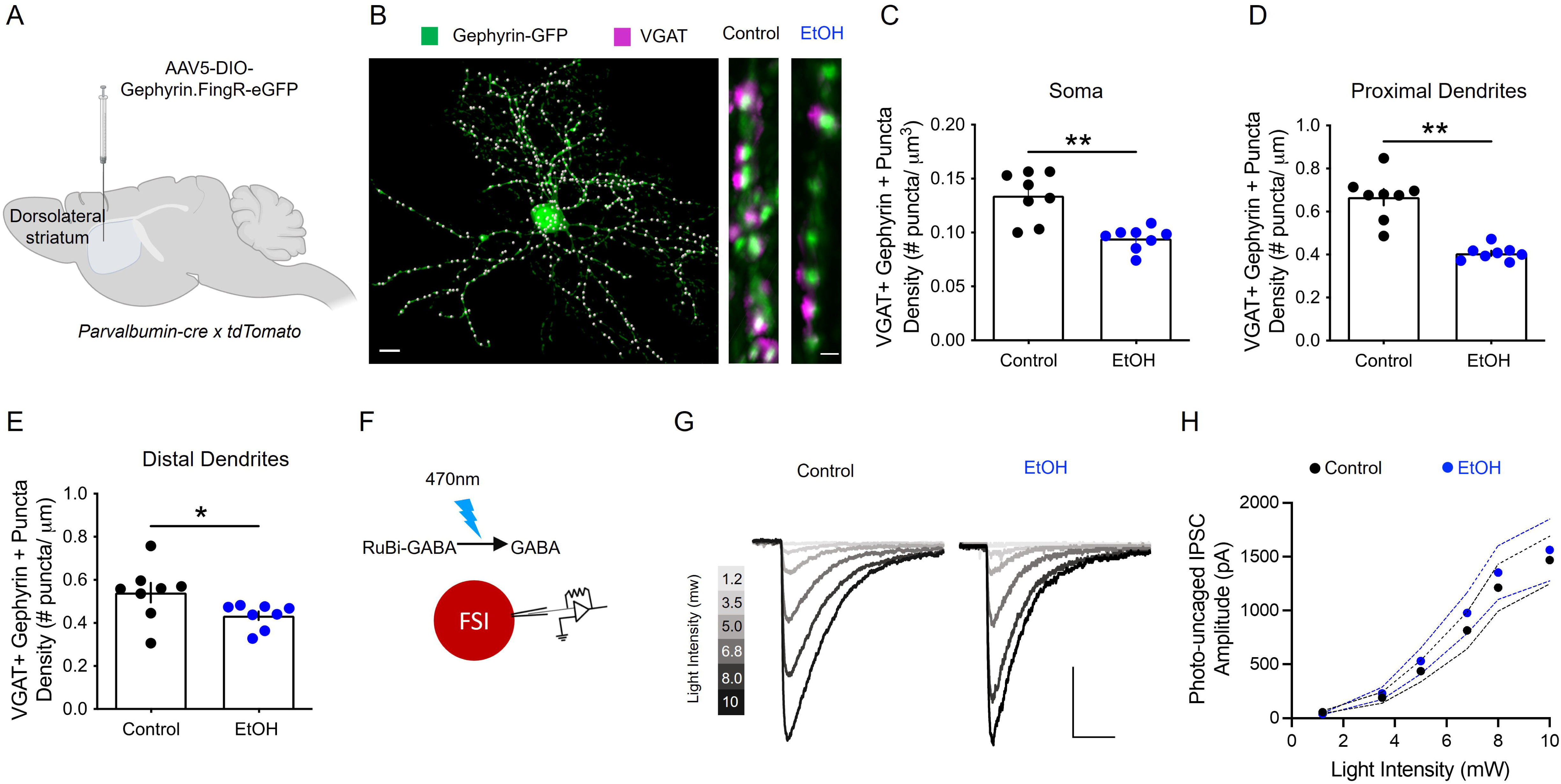
CIE de-localizes GABAergic synaptic contacts onto FSIs. **A**) Schematic of viral injection of virus expressing a cre-dependent Gephyrin-GFP intrabody into parvalbumin (PV)-cre mice. **B**) Representative image of an FSI expressing gephyrin-GFP intrabody (green) colocalized to an immunofluorescent marker for vesicular GABA transporter (VGAT) puncta. VGAT positive puncta were masked onto the GFP signal and colocalized signals (white dots) were labeled with a dot function model to quantify GABAergic synapses (scale bar 10µm). (right) Representative fluorescent image of proximal neuronal branch from control and CIE treated mice depicting colocalized presynaptic VGAT (purple) puncta onto gephyrin (green)-positive postsynaptic elements (scale bar 1µm). CIE dramatically reduced the density of colocalized VGAT puncta onto somata (**C**), proximal dendrites (**D**), and distal dendrites (**E**) (N=8 mice per group, 4M 4F). **F**) Schematic of whole-cell patch clamp recording from an FSI during photo-uncaging of RuBi-GABA. **G**) Representative traces of photo-uncaged GABA currents from FSIs of control (left) and CIE (right) treated mice at increasing stimulus intensities (scale bar = 200pA, 100ms). **H**) CIE did not impact photo-uncaged IPSCs compared to control treated mice (n=8 mice per group, 4M 4F). All data represented as mean + SEM. *P < .05, **p<.01. Portions of this figure were created with BioRender.com

As the reduction in putative GABAergic synapses was more prominent around the somata and proximal processes of FSIs, we hypothesized that a perisomatic mechanism accounts for cellular subregion-specific synaptic disruption. Because perineuronal nets (PNNs) condense around the somata and proximal branches of FSIs and are known to regulate synaptic transmission onto FSIs^42–44^, we investigated whether CIE modulated PNNs. We first confirmed that PNNs are indeed present in the dorsal striatum of adult C57BL/6J mice using *Wisteria floribunda* agglutinin (WFA) staining^45^. We identified WFA positive PNNs surrounding FSIs (**Figure 4A**), consistent with previous literature^46^. However, we noticed that some FSIs were enriched with PNNs (PNN+), and that a subpopulation were PNN poor (PNN-) based on a fluorescent signal intensity threshold for WFA staining (**Figure 4A-C**). We next examined the effect of CIE on PNN presence surrounding FSIs. CIE reduced the number of PNN+ FSIs (t_(14)_ = 12.83, p < 0.0001; Student’s t-test; **Figure 4D**), but had no impact on the total number of FSIs (t_(14)_ = 1.01, p = 0.33; Student’s t-test; **Figure 4E**). The reduction in PNN+ FSIs was not evident after 2-weeks of CIE (t_(10)_ = 0.85, p = 0.41; Student’s t-test; **S1A**), indicating that loss of PNN expression was specific to long-term ethanol exposure. We followed these results by western blotting for two PNN proteins; the proteoglycan aggrecan^22^, and the link protein hyaluronan and proteoglycan link protein 1 (HAPLN1)^44,47^. Consistent with our population analysis, aggrecan (t_(14)_ = 3.87, p < 0.01; Student’s t-test; **Figure 4F)** and HAPLN1 (t_(14)_ = 2.91, p < 0.05; Student’s t-test; **Figure 4G)** expression was reduced in dorsolateral striatum punches from CIE compared to control mice.

**Figure 4.**
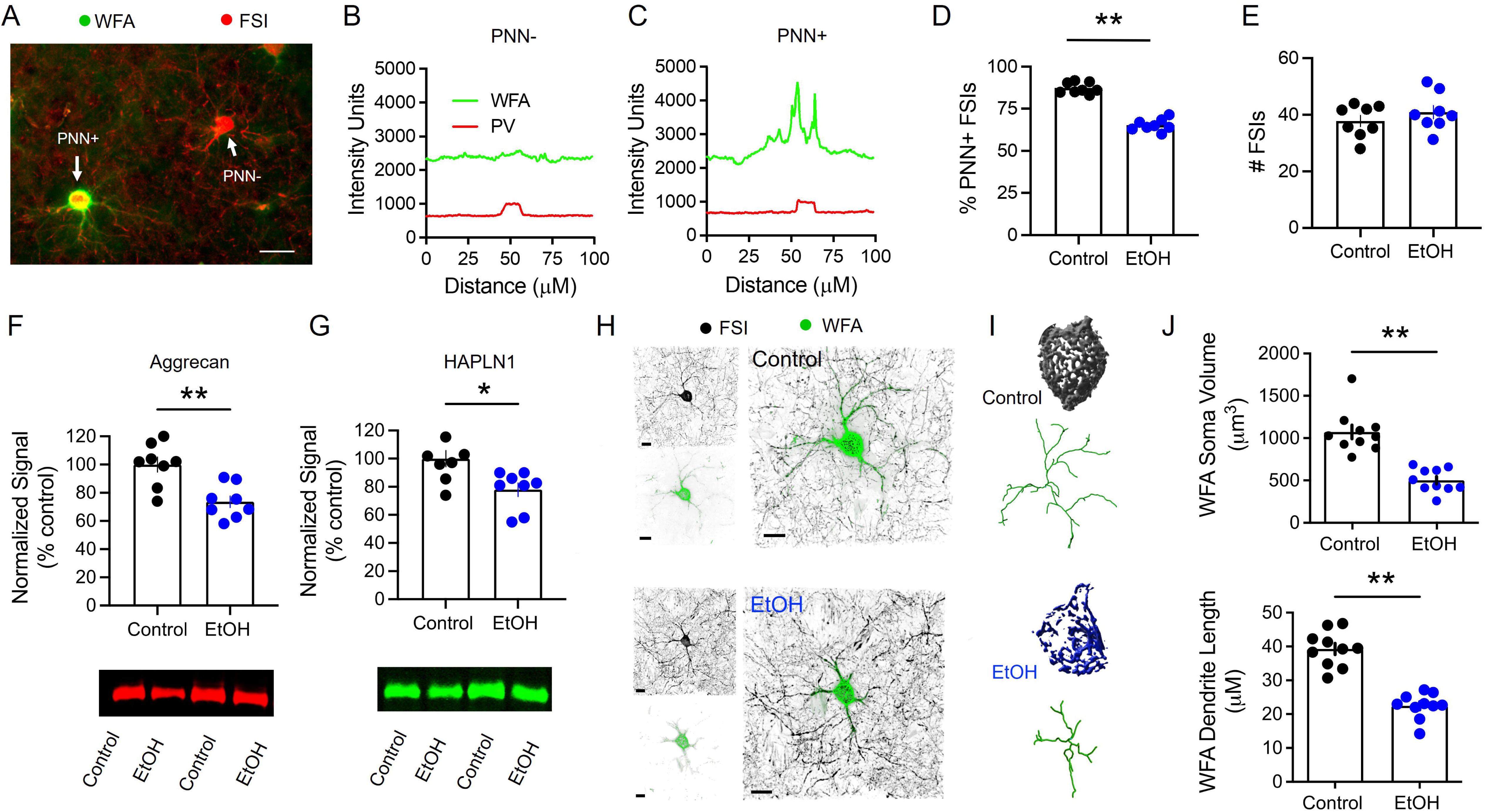
CIE degrades perineuronal nets (PNNs) in the dorsolateral striatum. **A**) Immunofluorescent image of parvalbumin+ FSIs (red) and WFA+ perineuronal nets (green) depicting net enriched (PNN+) and net poor (PNN-) FSIs (scale bar 100µm). **B,C**) Fluorescent intensity profile of PNN+ and PNN-FSIs. D) CIE reduced the percentage of net enriched FSIs but (**E**) does not reduce the number PV+ FSIs (N=10 mice per group, 5M 5F). CIE reduced the expression of PNN proteins (**F**) aggrecan and (**G**) HAPLN1 in the dorsolateral striatum (N=8 mice per group, 4M 4F). **H**) Representative confocal fluorescent images of PV+ FSI (black) and WFA+ PNN (green) from control (top) and CIE (bottom) treated mice (scale bar 15µm). **I**) Top: 3D volume reconstruction and filament model of WFA fluorescent signal from control (top) and CIE (bottom) treated mice. **J**) CIE reduced the volume of WFA signal (top) surrounding FSI somata and reduced the length of WFA signal (bottom) surrounding FSI proximal branches (N=10 mice per group, 5M 5F). All data represented as mean + SEM. *P < 0.05. ** P < 0.01.

To determine whether specific cellular sub-compartments were impacted by CIE, we collected confocal images of WFA PNN+ signal from control and CIE mice (**Figure 4H**) and generated 3D surface expression models surrounding FSI somata and branches (**Figure 4I**). Analysis of these models revealed a significant reduction in WFA signal surrounding FSI somata (t_(18)_ = 6.24, p < 0.0001; Student’s t-test; **Figure 4J**) and a reduction in the length of WFA signal extending along FSI branches (t_(18)_ = 8.23, p < 0.0001; Student’s t-test; **Figure 4J**). However, there was no difference in the branch length of the FSIs between groups (t_(18)_ = 0.03, p = 0.98; Student’s t-test; **S1B**).

During brain development, PNNs regulate the formation of synaptic connections^48^. We therefore investigated whether PNNs regulated GABAergic transmission onto FSIs in the dorsolateral striatum. We first recorded electrically-evoked GABAergic transmission from FSIs and later identified them as PNN+ or PNN-by dye-filling the recorded cell and subsequent WFA staining (**Figure 5A**). PNN+ FSIs exhibited larger electrically-evoked IPSC amplitudes compared to PNN-FSIs (F_(1,15)_ = 8.10, p < 0.05; two-way repeated measures ANOVA; **Figure 5B**). This difference was not due to a change in presynaptic release probability, as no mean difference in the PPR was observed (t_(15)_ = 0.74, p = 0.47; Student’s t-test; **Figure 5C**). Furthermore, PNN+ FSIs exhibited greater frequency of sIPSC events compared to PNN-FSIs (t_(13)_ = 2.53, p < 0.05; Student’s t-test; **Figure 5D, E**), but there was no change in sIPSC amplitude (t_(13)_ = 0.56, p = 0.58; Student’s t-test; **Figure 5F**).

**Figure 5.**
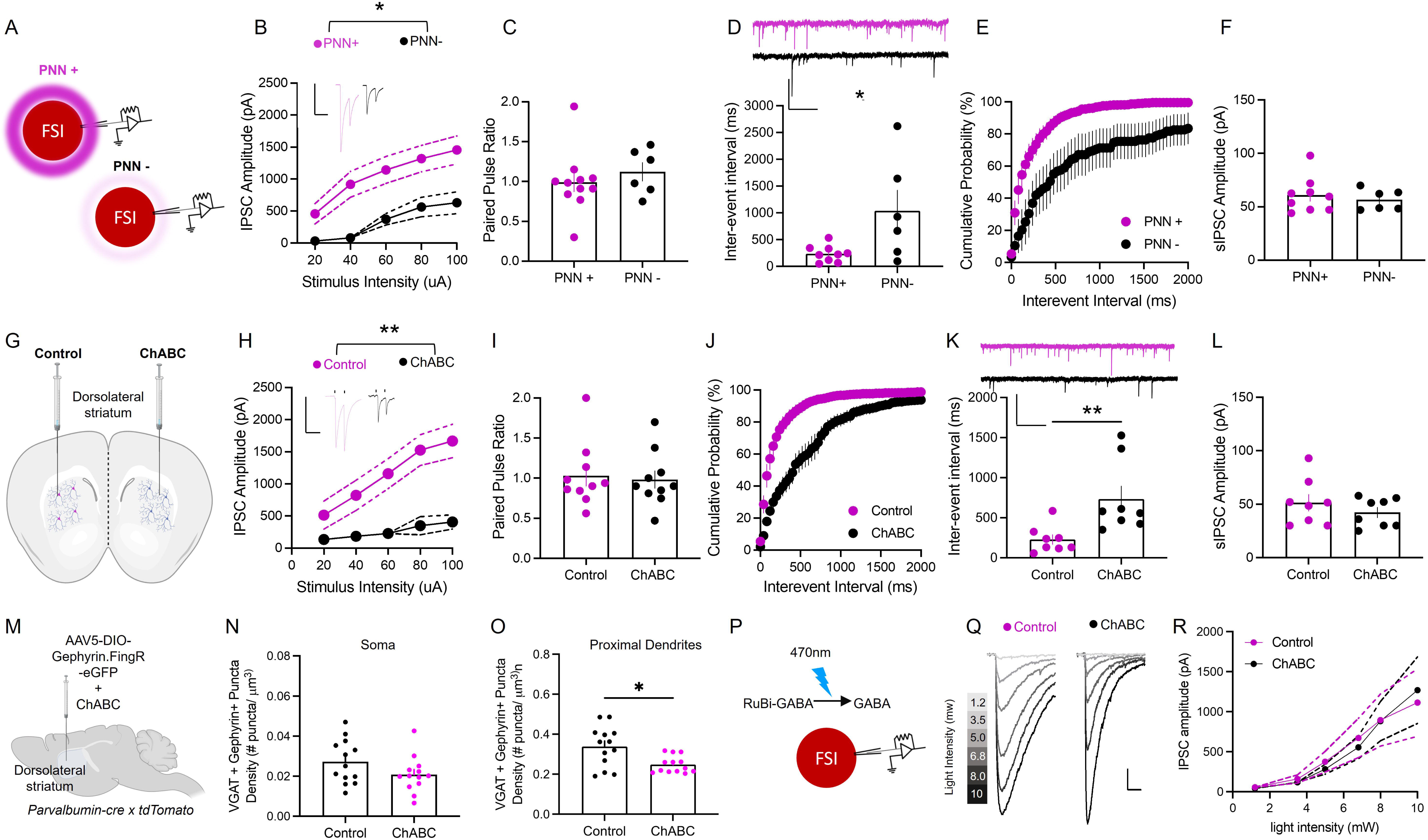
PNN enzymatic digestion suppresses GABAergic synaptic transmission onto FSIs. **A**) schematic of whole-cell patch recording from PNN enriched (PNN+) or PNN poor (PNN-) FSI identified by WFA stain after recordings. **B**) PNN+ FSIs exhibited larger eIPSC amplitudes compared to undetected PNN-FSIs (N=11 PNN+ cells, 5M 6F; 6 PNN-cells, 3M 3F). Inset: representative traces from PNN+ (pink) and PNN+ (black) FSIs (scale bar 1nA, 100ms). **C**) The presence of PNNs did not impact GABA release probability onto FSIs. **D-F**) Spontaneous IPSC events (sIPCSs) recorded from FSIs (N=11 PNN+ cells, 5M 6F; 6 PNN-cells, 3M 3F). Inset: representative traces from PNN+(pink) and PNN-(black) FSIs (scale bar 100pA, 1s). **D,E**) PNN+ FSIs exhibited a greater frequency of spontaneous IPSC events but (**F**) no change in IPSC amplitude compared to PNN-FSIs. **G**) Schematic of unilateral microinjection of chondroitinase ABC (ChABC) into the dorsolateral striatum to enzymatically degrade PNNs. **H**) Degrading PNNs with ChABC reduced eIPSC event amplitude onto FSIs but (**I**) did not impact GABA release probability (N=10 mice, 5M 5F; scale bar 1nA, 100ms). **J,K**) Degrading PNNs with ChABC reduced the frequency of sIPSC events onto FSIs but (**L**) did not impact event amplitude (N=8 mice, 4M 4F; scale bar 100pA, 1s). **M**) Schematic of viral injection of cre-dependent Gephyrin-GFP intrabody into PV-cre mice followed by ChABC. **N)** Degrading PNNs with ChABC did not reduce inhibitory synapses onto somata but (**O**) did reduce inhibitory synapse number on proximal dendrites (N= 13 mice, 6M 7F). **P**) Schematic of whole-cell patch clamp recording during photo-uncaging of RuBi-GABA onto FSIs treated with ChABC. **Q**) Representative traces of photo-uncaged GABA postsynaptic currents on control (left) and ChABC (right) treated FSIs at increasing stimulus intensity (scale bar = 200pA, 100ms). **G**) There was no change in photo-uncaged IPSC event amplitude between control (black) and ChABC (pink) FSIs (N=8 mice per group, 4M 4F). All data represented as mean ± SEM. *P < 0.05. ** P < 0.01. Portions of this figure were created with BioRender.com

We next examined whether PNN degradation causally decreased GABAergic transmission to FSIs. To do this, we enzymatically degraded PNNs with chondroitinase ABC (ChABC), which dramatically reduced the number of PNN+ FSIs at 3-4 days following stereotaxic microinjection into the dorsolateral striatum (F_(2,34)=_105, P<0.0001; Sidak’s multiple comparison test; **S2A,B**) compared to the VEH-injected control side. This effect was still evident at 14 days post injection (Sidak’s multiple comparison test; **S2B**). Notably, degrading PNNs had no impact on the number of FSIs (F_(2,34)=_1.9, p = 0.15; **S2C**). Next, we recorded from dorsolateral striatum FSIs and compared GABAergic transmission from control and ChABC treated hemispheres (**Figure 5G**). Enzymatically degrading PNNs reduced electrically-evoked IPSCs onto dorsolateral striatum FSIs (F_(1, 18)_ = 14.89, p < 0.01; two-way repeated measures ANOVA; **Figure 5H**). This reduction in GABAergic transmission was associated with a change in PPR (t_(18)_ = 0.27, p = 0.79; Student’s t-test; **Figure 5I**), but reduced the frequency of sIPSC events (t_(14)_ = 3.0, p < 0.01; Student’s t-test; **Figure 5JK**) with no impact on sIPSC event amplitude (t_(14)_ = 1.0, p = 0.33, Student’s t-test; **Figure 5L**).

We next determined if ChABC de-localized GABAergic synaptic contacts onto FSIs as we observed following CIE. Again, we virally expressed the GFP-labeled intrabody against gephyrin in dorsolateral striatum FSIs (**Figure 5M**). Following four weeks of viral expression, mice then received a unilateral microinjection of ChABC or 5% BSA vehicle control into the dorsolateral striatum. ChABC treatment reduced co-localized GABAergic synapses onto proximal dendrites (t_(12)_= 2.66, p < 0.05; Paired t-test; **Figure 5O**) but not somata (t_(12)_= 1.57, P = 0.14; Paired t-test; **Figure 5N**) of dorsolateral striatum FSIs compared to control mice. However, uIPSCs onto FSIs from control and ChABC treated mice were not significantly different, suggesting that postsynaptic inhibitory synapses remain unsilenced (F_(1,7)_ = 0.016, p = 0.9; **Figure 5Q,R**).

## Discussion

CIE specifically reduced GABAergic transmission onto dorsolateral striatum FSIs, including from the GP and the RTN, and modeling this effect by silencing GP and RTN inputs to the dorsolateral striatum increased alcohol drinking. Exploring the mechanism underlying GABA synapse de-localization, CIE degraded FSI PNNs, which we found was causal to GABA synapse de-localization and suppressed inhibitory control of FSIs. These data are consistent with a model wherein chronic alcohol exposure suppresses inhibitory control of dorsolateral striatum FSIs to promote alcohol drinking behavior through a PNN degradation-dependent process.

The dorsolateral striatum encodes habitual actions underlying compulsive behavior seen in substance use disorder^3, 4^. Dorsal striatum FSIs play a critical role in the expression of habitual behavior^18^, including compulsive-like ethanol consumption^13^. Here we demonstrate that chronic ethanol exposure reduces GABAergic, but not glutamatergic input onto dorsal striatum FSIs. In response to acute ethanol, FSI intrinsic excitability is increased^26^, GABA synaptic transmission onto FSIs is increased^23^, but GABA release from FSIs onto postsynaptic spiny projection neurons is decreased^25^. Provided the critical role of striatal FSIs for neuronal ensemble formation^49^, understanding the complex interactions between FSI acute and chronic alcohol exposure effects on the development of striatal ensembles, including those recruited for drug acquisition^20, 22, 50^, is positioned to provide deeper insight to the development of learned, inflexible behaviors in alcohol use disorder.

The effect of CIE on GABAergic transmission was evident in two major GABAergic projections the dorsolateral striatum, the GP and the RTN, suggesting a cell-wide modification of GABAergic input. Indeed, GABAergic transmission from GP and RTN inputs to FSIs is increased in response to acute ethanol exposure^23^. Whether the FSI-FSI synapse is affected by either acute or chronic alcohol exposure is yet to be investigated^51^. How multiple acute alcohol exposures, which by themselves which enhances GABA, give rise to the GABA synapse de-localization that suppresses inhibitory control of FSIs is a major question arising from the present findings. Given the evidence for PNNs in this process, multiple cellular mechanisms could underly this phenomenon^52^, including synaptic pruning^53^, possibly mediated by microglia^54–57^. Another possible mechanism is ethanol’s modulation of PNN-degrading enzymes released by neurons and glia, including metalloproeinase-9 (MMP-9) enzyme^44^. In humans, serum MMP-9 levels increase during alcohol intoxication^58^ and in individuals with a history of alcohol abuse^59^. Similarly, in rats, MMP-9 expression is elevated in the nuclues accumbens of alcohol dependent rats^60^. Notably, intracerebroventricular administration of a broad-spectrum MMP inhibitor reduces the escalation of ethanol self-administration in rats^61^.

PNNs condense around the soma and proximal processes of FSIs to stabilize synaptic contacts and regulate synaptic transmission^44, 62–64^. We demonstrate that chronic ethanol exposure decreases the number of FSIs containing PNNs and decreases PNN protein expression in the dorsolateral striatum. This result aligns with the ethanol-induced PNN loss reported in the retrosplenial cortex^65^ and hippocampus^66^. However, in the insular^67^ and prefrontal cortex^68^, repeated ethanol exposure increases PNN expression. Regional differences in PNN expression are also observed following cocaine exposure^69^, suggesting that local microenviroments differentially respond to drugs of abuse resulting in increased or decreased PNN expression. Determining the mediators of these regionals effects may reveal important biology underlying neurobiological responses to drug of abuse exposure and resulting maladaptive behavioral changes.

Drug-induced PNN modification is thought to re-open a critical period of synaptic plasticity enabeling the formation of drug memories and drug-associated behaviors^70, 71^. Mirorring what is observed following CIE-induced PNN degradation, we found that TeLC silencing of GABAergic transmisson from the GP and RTN increased voluntary ethanol consumption in mice. These data suggest that chronic ethanol expsoure may be reopening a critical period for dorsal striatal-mediated learning that may contribute to inflexible alcohol consumption. Whether the increase in alcohol consumption we observed herein is mediated by an inflexible drinking phenotype requires further investigation. Regardless, the discovery that chronic alcohol exposure supresses induces remodeling of inhibitory control of FSIs, which are required for aversion resistant alcohol consumption, suggests future therapeutic strategies aimed at FSI PNN remodeling could present a promising approach for treating alcohol use disorder.

## Acknowledgments

This work was supported by The National Institute on Alcohol Abuse and Alcoholism grants, R01AA028070 (B.N.M), R01AA024845 (B.N.M.) and F31AA029264 (M.S.P.).

## Competing Interests

The authors declare no competing interests.

## Author Contributions

M.S.P and B.N.M. conceived experiments. M.S.P., P.N.M., J.W.V, M.H.P., and M.H. performed experiments. M.S.P. and B.N.M. wrote the manuscript.

## Notes

### Competing Interest Statement

The authors have declared no competing interest.

