## Supplemental Figures 1 and 2 for "Perineuronal Net and Inhibitory Synapse Remodeling on Striatal Fast-spiking Interneurons by Chronic Alcohol Exposure"

**Supplementary information**

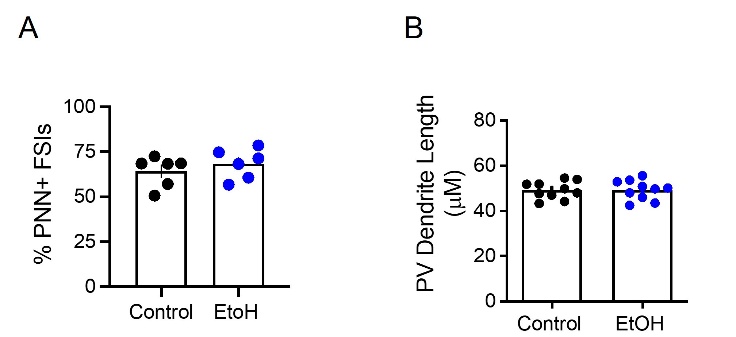

**Figure S1**. **Chronic intermittent ethanol (CIE) vapor exposure reduced perineuronal net (PNN) expression in the dorsolateral striatum**. **A**) Two weeks of CIE did not impact the number of PNN+ FSIs (N=6 mice per group, 3M 3F). **B**) Five weeks of CIE did not impact FSI dendrite length (N=10 mice per group, 5M 5F). All data represented as mean ± SEM.

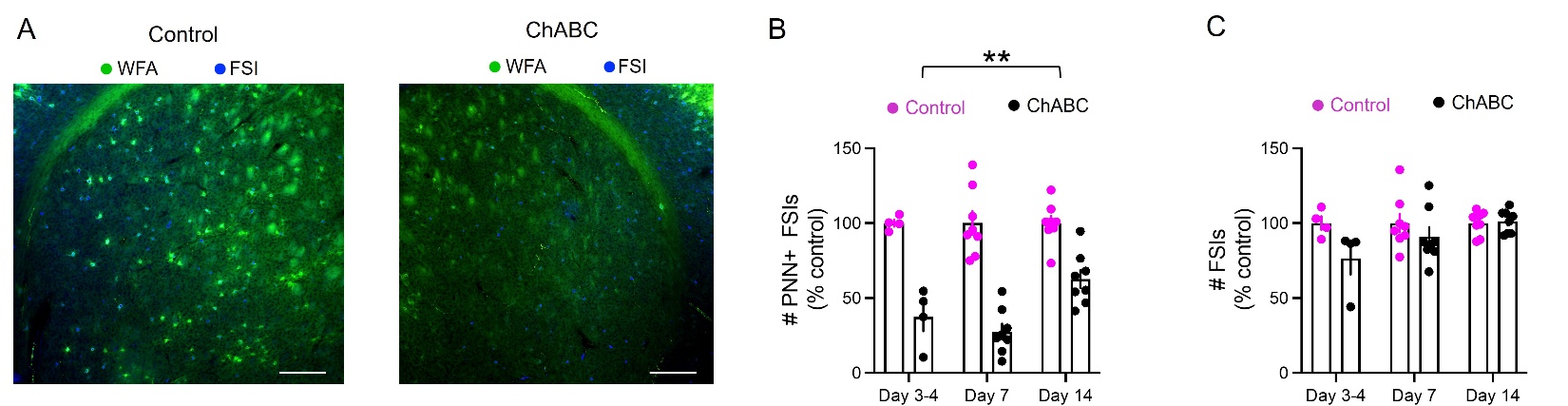

**Figure S2**. **Chondroitinase ABC (ChABC) reduced PNN expression in the dorsolateral striatum.** **A**) Representative images from control (left) and ChABC (right) microinjection into the dorsolateral striatum to degrade PNNs (green) surrounding FSIs (blue). Scale bar = 50µm.  **B**) Quantification of PNN+ FSIs 3-4 days, 7days, and 17 days following microinjection of ChABC or vehicle (VEH) control. ChABC reduced PNN+ FSIs in the dorsolateral striatum at least 3-4 days following microinjection and was still evident at 14 days post-microinjection. **C**) ChABC microinjection into the dorsolateral striatum did not impact the number of PV+ FSIs. All data represented as mean ± SEM. ** P < 0.01
